# How to control unpredictable others in court-net sports

**DOI:** 10.1101/2023.12.20.572699

**Authors:** Yuji Yamamoto, Keiko Yokoyama, Akifumi Kijima, Motoki Okumura, Hiroyuki Shima, Kazutoshi Gohara

## Abstract

Competition in sports does not involve a single solution because individuals aim to behave unpredictably, thereby preventing others from predicting their actions. This study determined how individuals in court net sports tried to control others’ unpredictable behavior, thereby addressing the gap of the lack of clarity about strategies employed by individuals in competitive situations. We employed a switching hybrid dynamics model, considering external inputs when analyzing individual behaviors. The study shows that skilled individuals, unlike intermediate players, exhibit greater regularity in their behavior, lead others to anticipate this regularity, and employ strategies to disrupt these expectations. This strategy exploits the principles of active human inference, implying that competition involves cooperation. We revealed this by analyzing both human decision-making and behavior in actual matches as discrete and continuous dynamical systems. This strategy helps policymakers adopt a new policy targeting cooperation with competitors, which would increase competitiveness in our daily lives.

## Introduction

Humans make decisions and take actions based on these decisions. Our decisions and actions influence others, and vice versa. In his retrospective paper (***Morgenstern, 1976***), the famous game theorist Morgenstern used the example of Professor Moriarty’s battle with Sherlock Holmes to illustrate a strategy in game theory. He refers to this strategy as a circular argument, involving arbitrary decisions such as “I think he thinks that I think!!…”. Interpersonal competition is similar to this circular argument and is a problem, such as an ill-posed problem, to which a unique solution cannot be determined. However, game theory mathematically analyzes the interdependence of decision-making in economic activities and does not involve actions, such as those of Holmes as cited by Morgenstern (***Morgenstern, 1976***), and has not yet fully addressed interpersonal competition involving individuals’ physical activities in their daily life.

Interpersonal competition, as analyzed in game theory, is as fundamental to daily human life as interpersonal cooperation. The study of joint action suggests that, for successful cooperation, reducing variability in individuals’ actions and increasing predictability of these actions is important (***Vesper et al., 2011; Sabu et al., 2020***). These are called coordination smoothers (***Vesper et al., 2010***) which aim to enhance coordination by modifying actions, such as exaggerating movements (***Goebl and Palmer, 2009***) or reducing action variability. Conversely, competitive situations increase unpredictability (***Glover and Dixon, 2017***). However, the competitive situation in this study involves an asymmetrical leader-follower relationship, rather than a symmetrical reciprocal relationship commonly observed in sports or everyday life. In a reciprocal relationship, it is conceivable that both individuals increase the variability of their actions, making them less easily predicted by others.

However, from the perspective of the free energy principle or active inference, individuals are also expected to reduce uncertainty about others’ actions and minimize surprise (***Friston et al., 2006; Friston, 2010***). However, it can be argued that if others become unpredictable owing to increased variability in their actions, predicting their actions corresponding to one’s own actions, especially in sequential and reciprocal interactions, will become challenging. In other words, increasing the variability of one’s own behavior will increase the uncertainty of others’ behavior, making predictions more difficult.

Strategies employed by individuals in both sequential and reciprocal competitive situations remain largely unclear. Therefore, this study aimed to elucidate how individuals attempt to control unpredictable others in competitive situations. To achieve this objective, this study focused on court net sports, which involves players hitting a ball alternately rather than simultaneously, competing against each other while their goals are in opposition. Players’ decision-making can be inferred from the direction of the ball’s trajectory, while their actions or responses to others can be observed as movements in court. Furthermore, quick decision-making and the execution of appropriate responses are required, and skilled players undergo extensive practice to improve, making them well-suited for observing refined strategies in sequential and reciprocal competitive situations.

Interpersonal competition in court-net sports requires successive decision-making and abrupt switching of movement patterns corresponding to unpredictable environmental changes, such as forehand and backhand strokes corresponding to opponents’ shots. In other words, to switch movement patterns, players must perceive environmental changes, such as opponents’ shots, and make decisions based on players’ intentions and states. To describe these phenomena, the dynamical system model should consider the switching movement corresponding to the external temporal input and discrete decision-making with continuous movement. We introduce a switching hybrid dynamical system (SHDS) model (***Nishikawa and Gohara, 2008a***,b) to satisfy these requirements.

The SHDS model comprises a discrete dynamical system with an external temporal input and a continuous dynamical system connected to a feedback loop. Discrete and continuous dynamical systems correspond to the brain and body, respectively, (***Yamamoto et al., 2018***). In the discrete dynamical system, as a higher module, decisions are made corresponding to the external temporal input, and these decisions are sent to the continuous dynamical system. In the lower module, the continuous dynamical system switches the movement pattern corresponding to decisions from the discrete dynamical system. The output pattern of the continuous dynamical system was fed into a discrete dynamical system as a feedback signal. Consequently, in the discrete dynamical system the next decision is made based on both the current external temporal input and its own physical state. If the external temporal input was constant and repetitive, the movement pattern would converge in a stable state as an excited attractor. However, if the external temporal input is switched in some time intervals, the movement pattern also switches corresponding to the external input; thus, the movement pattern shows fractal transitions between excited attractors, and we can observe hysteresis (***Gohara and Okuyama, 1999a***,b) (see detail ***Appendix 1***).

This means that the discrete dynamical system corresponds to decision-making depending on opponent’s’ shots and players’ own movements or preparatory stances, and the continuous dynamical system corresponds to the pattern of players’ hitting movements. A switching dynamics model with an external temporal input was examined using striking movements under well-controlled experimental conditions. These experiments introduced a pseudo-random sequence of temporal inputs, that is, a third-order sequential effect (***Remington, 1969; Kirby, 1976; Soetens et al., 1984***), and the fractal transitions between two excited attractors were confirmed, that is, the hysteresis for forehand and backhand strokes in tennis (***Yamamoto and Gohara, 2000***) and the fractal dimensions in table tennis (***Suzuki and Yamamoto, 2015***). However, in these experiments, the temporal input was controlled experimentally, unlike in real games, and because they only observed individual movement and not interpersonal competition, individual decision-making was not required in the experiments that only responded to the external input. In this study, the SHDS model was first applied to interpersonal competition in real matches as a two-coupled SHDS model (***Yamamoto et al., 2018***).

In this study, we first confirmed that court-net sports can be described as a two-coupled SHDS based on the SHDS model. Specifically, we represent the regularity of the sequences of shot angles for two players in an actual game as a discrete dynamical system, and we represent the corresponding movement on court as a continuous dynamical system. The following hypotheses were then tested by examining the characteristics of the observed regularity in actual matches through numerical simulations of switching dynamics. Hypothesis: If individuals’ strategy is to increase the variability of their actions to make them less predictable to others, the sequence of shot angles will be closer to a more random sequence or if their strategy is to decrease the variability of their own actions and increase the predictability of others, ultimately increasing their own predictability, the sequence of shot angles will become closer to a regular (alternating left-right) sequence.

## Results

### Analysis of real matches

#### Regularity in a shot angle sequence as discrete dynamics

We confirmed the skill difference between international and collegiate matches in rally length (***Appendix 2***) and then examined regularities in a sequence of shot angles. We focused only on long rallies with more than nine successive shot sequences. To examine the regularity in the sequence of the shot angle Θ and shot length ΔL, return map analysis was applied to the data at both levels. Based on *R*^2^ and the significance of the regression model, we counted the number of sequences that were significantly fitted to the linear function *X*_*n*+1_ = *a X*_*n*_ +*b*. ***Figure 1*** shows the frequencies of well fitted results for the sequences including four to eight successive shots which were calculated from the data of the actual match and artificial surrogate data. In international matches, significant differences in the ratio of the significantly fitted shot angle Θ sequence between actual and artificial surrogate data were found in sequences, including four to eight successive shots (4 shots: *χ*^2^(1) = 5.95, *p* = 1.47 × 10^−2^, 5 shots: *χ*^2^(1) = 41.74, *p* = 1.04 × 10^−10^, 6 shots: *χ*^2^(1) = 52.86, *p* = 3.58 × 10^−13^, 7 shots: *χ*^2^(1) = 69.71, *p* < 2.20 × 10^−16^, 8 shots: *χ*^2^(1) = 150.18, *p* < 2.20 × 10^−16^), However, a significant difference between the actual and artificial data in the ratio of the significantly fitted shot length ΔL sequences was found only in the rally including four and six successive shots (four shots: *χ*^2^(1) = 9.42, *p* = 2.15 × 10^−3^; six shots: *χ*^2^(1) = 6.64, *p* = 9.97 × 10^−3^). On the other hand in collegiate matches, significant difference for the shot angles Θ was found in four and five successive shots (4 shots: *χ*^2^(1) = 17.24, *p* = 3.29 × 10^−5^, 5 shots: *χ*^2^(1) = 6.98, *p* = 8.26 × 10^−3^), and for the shot length ΔL sequences were found only in four successive shots (*χ*^2^(1) = 5.83, *p* = 1.57 × 10^−2^).

**Figure 1.**
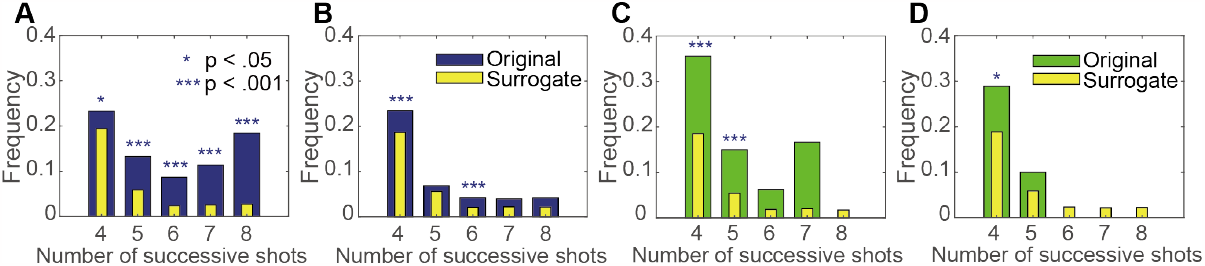
Frequencies of significant fitting for match data and surrogate data. A and B show the frequencies of the shot angle Θ and the shot length ΔL for the international match, respectively. C and D show the frequencies of the shot angle Θ and the shot length ΔL for the collegiate match, respectively. Blue and green bars show the frequencies of the significant fitting for the original data and yellow bars show the frequencies of significant fitting for the surrogate data. For a full definition of “international” and “collegiate” matches corresponding to expert and intermediate skill level respectively, please see the methods section. **Figure 1—figure supplement 1**. Means and standard deviations (SDs) of the fitting ratio by the number of random shuffling repetitions for surrogate data. A shows the change in the mean of the fitting ratio for 20—200 repetitions. B shows the change in SD of the fitting ratio for 20—200 repetitions.

Thus, in international matches, the significance of the shot sequence in fitting the linear functions was found in the sequence with all to 4–8 shots. Additionally, such significance to fit was found only in the analysis of shot angle Θ and not in the analysis of shot length ΔL. Moreover, the significance of the sequence of shot angles Θ was found in the data measured in both international and collegiate matches. Therefore, shot angle Θ is a more appropriate state variable that represents regularity in a rally for a real match. In particular, this regularity would strongly constrain longer sequences but not short sequences. Therefore, we focus on the shot angle Θ as the state variable of a discrete dynamical system and present the results of the analysis concerning the shot angle Θ only.

***Table 1*** shows each number (frequency ratio) of the sequence of the shot angle significantly fitted to the each of four functions, such as such as those of a rotational repeller, rotational attractor, asymptotic repeller, and asymptotic attractor. The sum of well-fitted data for both functions was 288 for international matches and 38 for collegiate matches. The significance of the ratio of data significantly fitted to each of the four functions was tested through their comparison with each ratio calculated from the surrogate data. The results revealed that the ratio calculated from the actual data was significantly higher for a rotational repeller and rotational attractor. However, for an asymptotic repeller and asymptotic attractor, only 16 and 12 cases in international matches, respectively, were significantly fitted to the function. Thus, in each of the international and collegiate matches, 90.3 % and 86.8 % of the significantly fitted sequences fluctuated in periodic patterns, respectively. Therefore, international players alternately changed their shot angles on the right and left sides against the opponent preceding the shot angle.

**Table 1.** Frequencies of significant fitting in original data in each type of attractor and repeller.

| Shots | International |  |  |  |  |  | Collegiate |  |  |  |  |  |
| --- | --- | --- | --- | --- | --- | --- | --- | --- | --- | --- | --- | --- |
|  | R<sub>rot</sub> | A<sub>rot</sub> | R<sub>asy</sub> | A<sub>asy</sub> | Sum | N | R<sub>rot</sub> | A<sub>rot</sub> | R<sub>asy</sub> | A<sub>asy</sub> | Sum | N |
| 4 | 71 | 53 | 14 | 8 | 146 | 626 | 17 | 11 | 3 | 1 | 32 | 90 |
|  | .486 | .363 | .096 | .055 | .233 |  | .531 | .344 | .094 | .031 | .294 |  |
| 5 | 31 | 22 | 1 | 4 | 58 | 479 | 5 | - | 1 | - | 6 | 40 |
|  | .534 | .379 | .017 | .069 | .132 |  | .833 | - | .167 | - | .150 |  |
| 6 | 13 | 13 | 1 | - | 27 | 343 |  |  |  |  |  |  |
|  | .481 | .481 | .037 | - | .087 |  |  |  |  |  |  |  |
| 7 | 9 | 17 | - | - | 26 | 252 |  |  |  |  |  |  |
|  | .346 | .654 | - | - | .114 |  |  |  |  |  |  |  |
| 8 | 5 | 26 | - | - | 31 | 187 |  |  |  |  |  |  |
|  | .161 | .839 | - | - | .184 |  |  |  |  |  |  |  |
| Total | 129 | 131 | 16 | 12 | 288 | 1770 | 22 | 11 | 4 | 1 | 38 | 130 |
|  | .448 | .455 | .056 | .042 | .163 |  | .579 | .289 | .105 | .026 | .292 |  |

R<sub>rot</sub>: Rotational Repeller. A<sub>rot</sub>: Rotational Attractor. R<sub>asy</sub>: Asymptotic Repeller.
A<sub>asy</sub>: Asymptotic Attractor. N: Number of all cases for fitting analysis.
Ratio of fitted number against the sum of fitted data. Ratio of fitted number against N.

***Figure 2*** A-D shows the typical examples of the time series of five successive shot angles Θ that largely and periodically (or rotationally) fluctuated (panel A and B) or slightly (or asymptotically) diverged or converged (panel C and D). Each ***Figure 2*** E-H shows a return map constructed from the data shown in ***Figure 2*** A–D. The time series shown in ***Figure 2*** A and C apparently differ in the amplitude and pattern of fluctuation (i.e., periodically or asymptotically); however, as can be seen in ***Figure 2*** E and G, both dynamical patterns can be similarly categorized as repellers on the return map. Such a similarity in the dynamical order between different time-series patterns can also be confirmed in ***Figure 2*** F and H, which are both categorized as attractors. The observed regularities for each rally per player at both levels are shown in ***Figure 2—figure Supplement 1*** and ***Figure 2—figure Supplement 2***.

**Figure 2.**
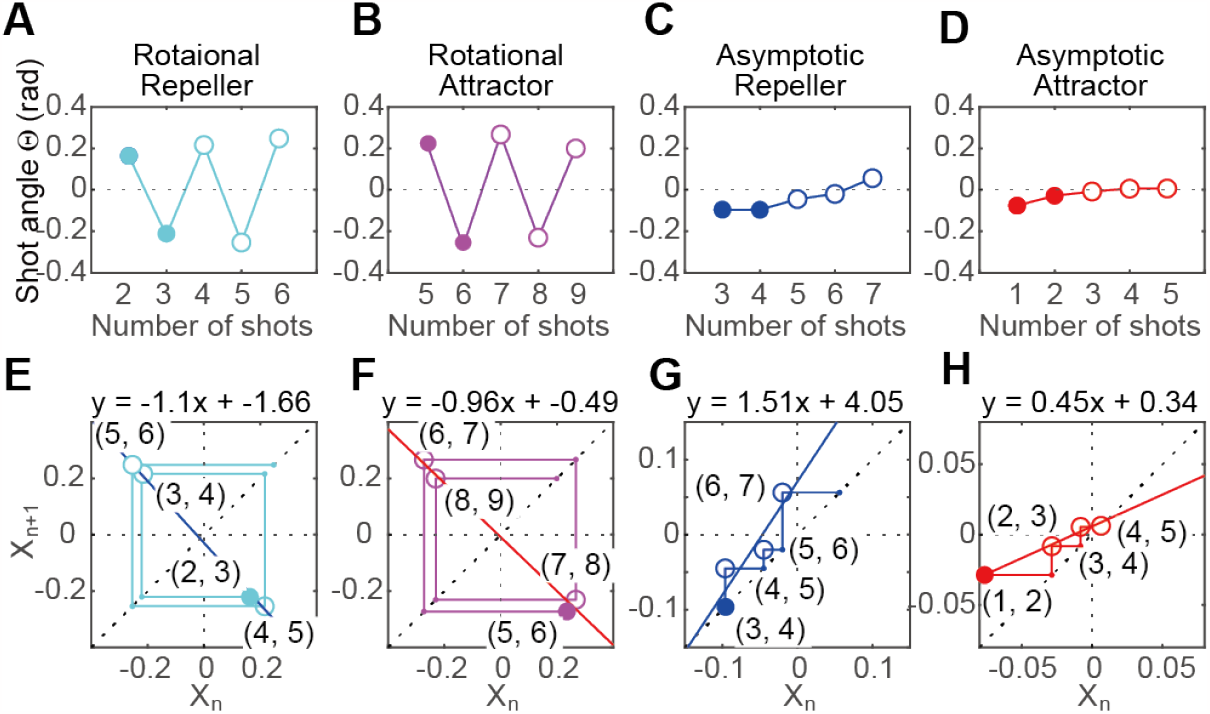
Typical examples of four types of attractors and repellers. A-D shows time series of the rotational repeller, rotational attractor, asymptotic repeller, and asymptotic attractor, respectively. E-H shows return maps corresponding to the above time series. **Figure 2—figure supplement 1**. Observed regularities in each rally per player in international matches **Figure 2—figure supplement 2**. Observed regularities in each rally per player in collegiate matches

The rotational-repeller (attractor) represents such dynamics, in which the shot angle gradually goes far (near) against the opponent’s previous shot, while changing the shot course alternatively to the right and left sides against the opponent’s preceding shot angle. In a relatively short sequence, which included five successive shots, the rotational-repeller appeared more frequently than the rotational attractor did. However, in longer sequences, including more than seven successive shots, attractors appeared more frequently than repellers did. These results suggest that the shot angles gradually increased as the total number of shots increased to 10—12 (5—6 shots x 2 players), and then the shot angle gradually decreased as the rally continued while both players kept changing their shot course referring to the course of the opponent’s shot. This means that the successive decision-making of the international player had the regularity of switching the course between the right and left sides in their longer rally.

#### Hysteresis on the continuous hitting movements

Second-order state transition probabilities, consisting of the right or left sides, were calculated for a well-fitted sequence of shots as the external temporal input of discrete dynamics. This means that the two cases, including the preceding and current inputs, were the same—right-right (RR) or left-left (LL)—and the preceding and current inputs were different—left-right (LR) or right-left (RL). The output pattern corresponding to the external temporal input consisted of two state variables.

In other words, polar angle *ϕ* and tangential velocity *v*_*ϕ*_ were depicted in the hyper cylindrical phase space ℳ as trajectories starting at the moment the opponent hit a ball, i.e., the Poincaré section Σ(*θ* = 0), until the opponent hits the next ball, i.e., Σ(*θ* = 2*π*) as continuous dynamics as the lower module. The trajectory around the cylinder represented one cycle from the current opponent’s shot to the next opponent’s shot, including one shot in the cycle. The first column in ***Figure 3*** A shows the state transition probabilities as a discrete dynamical system and the output pattern as a continuous dynamical system in all cases for international matches. The second and third columns show separate patterns for the right and left sides, respectively, as the current input. The trajectories corresponding to the right and left inputs were considered separately. ***Figure 3*** B shows the mean trajectories corresponding to the four kinds of external temporal input by second-order sequence effect as discrete dynamics. To provide a more detailed observation, the distributions of the Poincaré maps on the five Poincaré sections Σ, that is, *θ* = 0, *π/*2, *π*, 3*π/*2, and 2*π* are shown in ***Figure 3*** C. ***Figure 3—figure Supplement 1*** shows the results of collegiate matches corresponding to ***Figure 3***.

**Figure 3.**
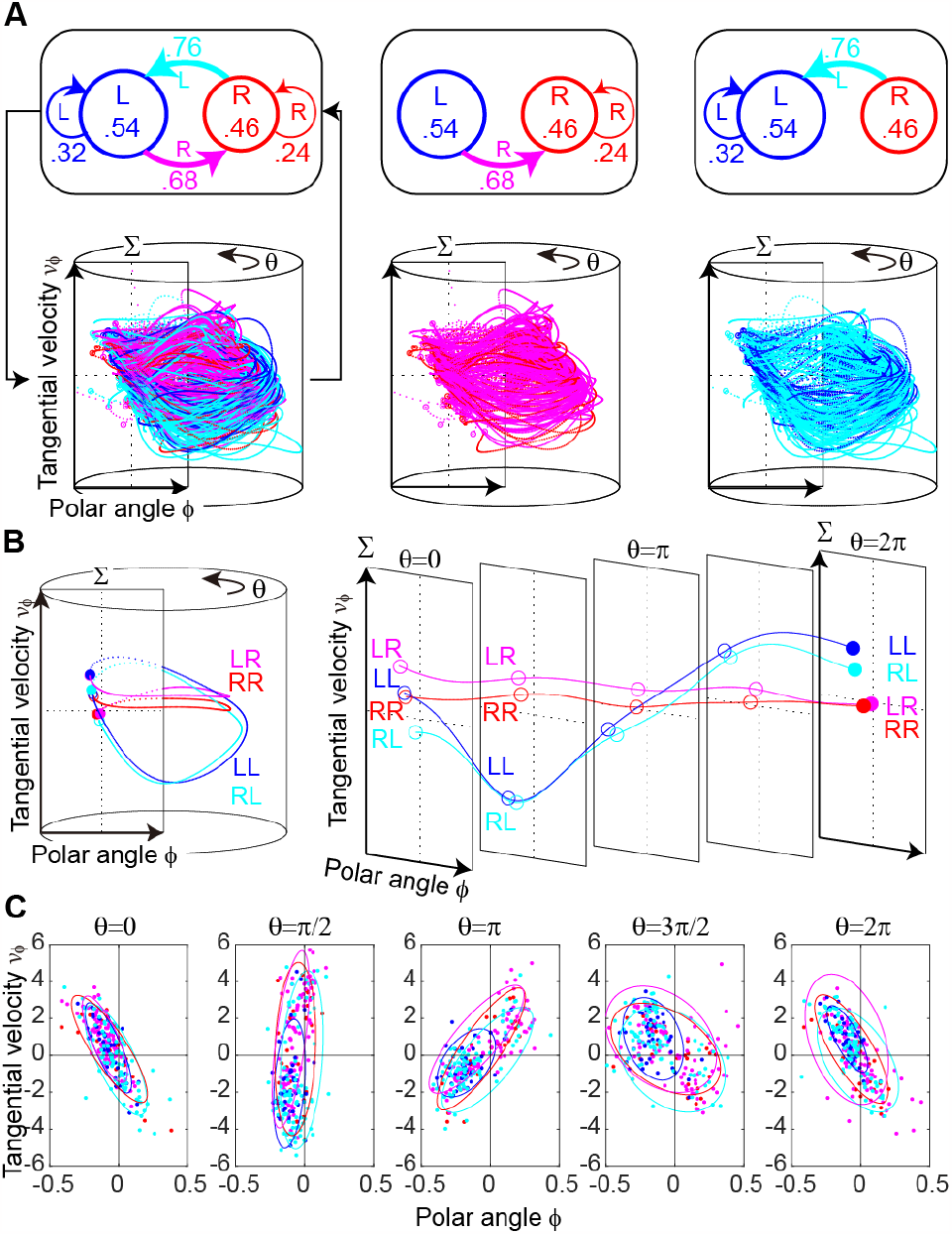
Switching hybrid dynamics in an international match. A show switching hybrid system in which discrete transition probabilities as a higher module are connected to continuous trajectories in a cylindrical phase space as a lower module. B shows that the mean trajectories are expanded into the two-dimensional plane, *θ* − (*x*_1_, *x*_2_) for switching input corresponding to panel a. The trajectories are separated corresponding to third-order sequence effects. C shows five Poincaré sections Σ, i.e., *θ* = 0, *π/*2, *π*, 3*π/*2, and 2*π* corresponding to panel B. with .8 equal probability ellipse. **Figure 3—figure supplement 1**. Switching hybrid dynamics in the collegiate match. A-C shows the same in ***Figure 3***. **Figure 3—figure supplement 2**. Hierarchical fractal structure. This demonstrates how the Cantor set can be constructed using two iterative functions. A and C show the iterative function *g*_*R*_ and *g*_*L*_ transform state *x*_*τ*_ to the next state *x*_*τ*+1_. In A, the lower-left and upper-right corners are the attractive fixed points 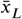 for *g*_*L*_ and 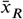 for *g*_*R*_, respectively. B and D show the hierarchical structure of the fractal corresponding to the sequence of right (R) and left (L) side inputs from *x*_0_ to *x*_3_. C and D are the same as A and B, except that the transformations of iterative function *g*_*R*_ and *g*_*L*_ are rotated around the fixed points 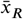 and 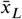, respectively. We named this the Cantor set with rotation. **Figure 3—figure supplement 3**. Eight trajectories of transitions between two excited attractors. The mean trajectories are expanded into the two-dimensional plane, using *θ* − (*x*_1_, *x*_2_) as the switching input. The trajectories are drawn with the third-order sequence effect. A shows eight mean trajectories. B and C show four mean trajectories corresponding to the right and left side inputs, respectively.

One-way MANOVAs were applied to test the equality of the multivariate means of the second-order sequence effect of the right and left inputs on each of the five Poincaré sections. As a result, in two Poincaré sections of five Poincaré sections, i.e., *θ* = 0 and *θ* = *π/*2, there were significant differences between Poincaré maps for the second-order sequence effects for international matches only (*θ* = 0: Preceding, Wilks Λ = .921, *F* (2, 251) = 10.782, *p* = 3.22 × 10^−5^, 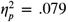; Current, Wilks Λ = .976, *F* (2, 251) = 3.147, *p* = .044, 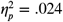, *θ* = *π/*2: Preceding, Wilks Λ = .935, *F* (2, 251) = 8.659, *p* = 2.31 × 10^−4^, 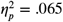; Current, Wilks Λ = .883, *F* (2, 251) = 16.637, *p* = 1.64 × 10^−7^, 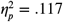). This result suggests that the spreading distributions in the Poincaré section did not result from random errors but depended on the sequence of the external temporal input. Namely, the output movement patterns depend not only on the current input but also on the preceding input; thus, we observed four clusters in Poincaré sections. This indicates that a hierarchical input sequence structure is mapped onto the excited attractor, and that the third-order sequence effect can be observed as trajectories in a hypercylindrical phase space(***Yamamoto and Gohara, 2000***).

We analyzed the characteristic configurations of the four clusters in Poincaré section Σ(*θ* = *π/*2) in ***Figure 3*** C. For example, the set of iterative functions *g*_*R*_ and *g*_*L*_ that provide a simple fractal structure (i.e., the Cantor set) was examined (see ***Figure 3—figure Supplement 2*** A). This simple example was introduced to demonstrate the behavior of dynamical systems excited by an external temporal input(***Gohara and Okuyama, 1999a***). This shows how the clustered structure corresponding to an input sequence is constructed from a set of iterative functions *g*_*R*_ and *g*_*L*_. The four clusters in the Poincaré section in the second stage *x*_2_ line up in the order of RR, LR, RL, and LL from the top (***Figure 3—figure Supplement 2*** B). This differs from the results of LR, RR, LL, and RL in Poincaré section Σ (*θ* = *π/*2) shown in Fig. 3 B. The sets of iterative functions (*g*_*R*_ and *g*_*L*_) in ***Figure 3—figure Supplement 2*** A were modified to provide those in ***Figure 3—figure Supplement 2*** C, with each iterative function having a rotatory attractive fixed point. Although they construct the same Cantor set, the sets in ***Figure 3—figure Supplement 2*** B and D provide significantly different cluster distributions. To emphasize this difference, the case in ***Figure 3—figure Supplement 2*** C is called a Cantor set with rotation. It is conjectured that the Cantor set with rotation originates from Hamiltonian dynamics with complex eigenvalues. The results of this study correspond to the Cantor set with rotation. The locations of the four clusters in the Poincaré section Σ (*θ* = *π/*2) (LR, RR, LL, and RL from the top), which are clearly second-order sequence effects, correspond to this theoretical prediction. Consequently, the eight clusters of trajectories were observed in the hypercylindrical phase space, as shown schematically in ***Figure 3—figure Supplement 3***. In summary, four clusters could be distinguished in the Poincaré section Σ and second-order sequence effects were confirmed. Further, there were eight trajectories between the excited attractors and third-order sequence effects in the hypercylindrical phase space. These sequence effects correspond to the hierarchical structure represented by the Cantor set with rotation.

These fractal transitions corresponding to the switching input provide strong evidence to support the SHDS. Furthermore, the switching input was not a well-controlled experimental condition (***Yamamoto and Gohara, 2000; Suzuki and Yamamoto, 2015***), which was the sequence of the opponent’s shot angle. This result supports our hypothesis that the individual dynamics underlying court-net sports can be understood as SHDS in a non-autonomous system.

#### Two-coupled switching hybrid dynamics

We found that 91 sequences in 60 rallies and 27 sequences in 19 rallies for international and collegiate matches, respectively, exhibited regularities in successive shot angles as a discrete dynamical system (***Figure 2—figure Supplement 1, Figure 2—figure Supplement 2***). Finally, we present a two-coupled SHDS in ***Figure 4*** as an example of sequences in which regularities are found in both players in a rally. The upper panels ***Figure 4*** A and A’ show the sequence of shot angles for both players as video clips and time series; ***Figure 4*** A shows for player X, and ***Figure 4*** A’ shows for player Y. ***Figure 4*** B and B’ show the return map corresponding to the sequence of ***Figure 4*** A and ***Figure 4*** A’, respectively. These systems are regarded as discrete dynamical systems with higher modules. The sequence of shots for player X (***Figure 4*** A) showed alternating changes between the left and right sides from the second shot to 8th shot. This sequence of shots for Player X from the second shot to 7th shot fitted the rotational repeller, as shown in ***Figure 4*** B. Conversely, the sequence of shots for player Y from 4th shot to 7th shot fitted the asymptotic repeller (***Figure 4*** B’). The sequence of shots for player Y, that is, the output of continuous dynamical systems for player Y (***Figure 4*** C’-F’), corresponds to the external temporal input *I*_*ext*_(*t*) for player X (***Figure 4*** A’ and B’) as the discrete dynamical system, and vice versa.

**Figure 4.**
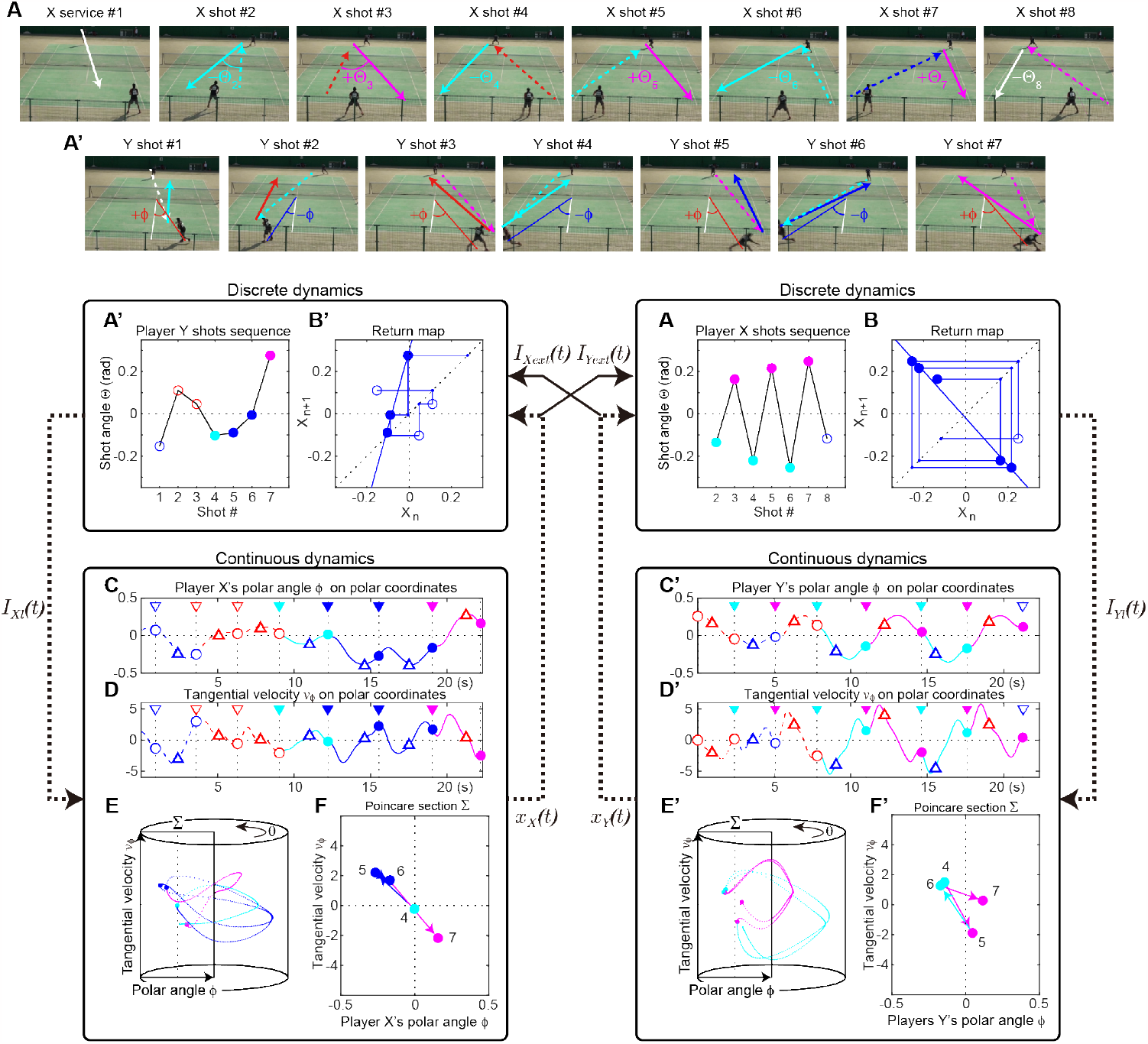
Two-coupled switching hybrid dynamics. Upper panel shows the sequences of both players’ shot angles in video clips in which the shot A’ show the time series of shot sequences corresponding to upper video clips. B and B’ show the return maps of A and A’. Blue circles show that the current input and the preceding input were the left side, and cyan circles shows the right side for the preceding input. Red circles show that the current input and the preceding input les also show the right side input but shows the left side for the preceding input. C and C’ show the time series yers in polar coordinates. D and D’ show the time series of tangential velocity of both players. The triangles show hot, and the circles show the moment of the players’ own shot. E and E’ show the trajectories in a cylindrical correspond to the four types of input. F and F’ show the Poincaré sections and the sequences of Poincaré maps. **Figure 4—figure supplement 1**. Partial correlation. A shows the partial correlation between the shot angle Θ and polar angle *ϕ*. B shows the hot angle Θ and tangential velocity *v*_*ϕ*_.

***Figure 4*** C and C’ show the time series of the players’ position on polar coordinates as angle *ϕ*, ***Figure 4*** D and D’ show the time series of tangential velocity as *v*_*ϕ*_. The inverse triangles show the moment of the opponent’s shot; colors show the direction—the right side (red) or left side (blue); and the triangles show the moment of their own shot. The positions of the shot corresponded to the external temporal input; when the input was on the right side, the shot position was on the right side (positive value), and vice versa. This means that the continuous dynamical system shows movements corresponding to the external temporal input; when the right-side input is fed into the system, the player moves to the right side to hit the ball and returns to the next waiting position. ***Figure 4*** E and E’ show the trajectories in a hyper cylindrical phase space consisted of two state variables, i.e., polar angle *ϕ* and tangential velocity *v*_*ϕ*_, and ***Figure 4*** F and F’ show the transitions of Poincaré map on the Poincaré sections Σ(*θ* = 2*π*).

To examine the relationship between the opponent’s shot course as an external temporal input and movement pattern as an output, partial correlation coefficients were calculated for international matches. The partial correlation coefficients between the shot angle Θ, polar angle *ϕ* and tangential velocity *v*_*ϕ*_ are 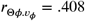, *p* = 1.31 × 10^−8^, 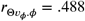, *p* = 4.92 × 10^−12^, respectively (***Figure 4—figure Supplement 1***). This suggests that the relationship between inputs and outputs is significant; namely, outputs strongly depend on inputs. The results showed that the two systems were connected both strongly and reciprocally, supporting our hypothesis that the dynamics of interpersonal competition underlying court-net sports could be considered as a two-coupled SHDS autonomous system.

### Simulation experiment

Players’ movements in a court in response to the continuous switching of external inputs were represented by a mass-spring-damper model. To investigate the regularity observed in a discrete dynamical system during real matches, the correlation dimensions on the Poincare map were calculated by modifying the second-order transition probabilities as a sequence of external temporal inputs. ***Figure 5*** A shows the correlation dimension on the Poincare map when the R-R and L-L state transition probabilities are independently modified. With perfect regularity, the R-R transition probability was 0, and the R-L transition probability was 1.0. Similarly, the L-L transition probability was 0 and the L-R transition probability was 1.0, resulting in an alternating repetition of different inputs. Complete regularity is located outside the lower-left quadrant of ***Figure 5*** A, where the corresponding Poincaré map is ***Figure 5*** E and the correlation dimension is zero. However, in the case of complete randomness, the R-R transition probability was 0.5; the R-L transition probability was 0.5; the L-L transition probability was 0.5; and the L-R transition probability was 0.5. This configuration is located at the center of ***Figure 5*** A, where the corresponding Poincaré map is shown in ***Figure 5*** B. At least eight clusters are observed, indicating the presence of a fractal structure with a correlation dimension of 1.433.

**Figure 5.**
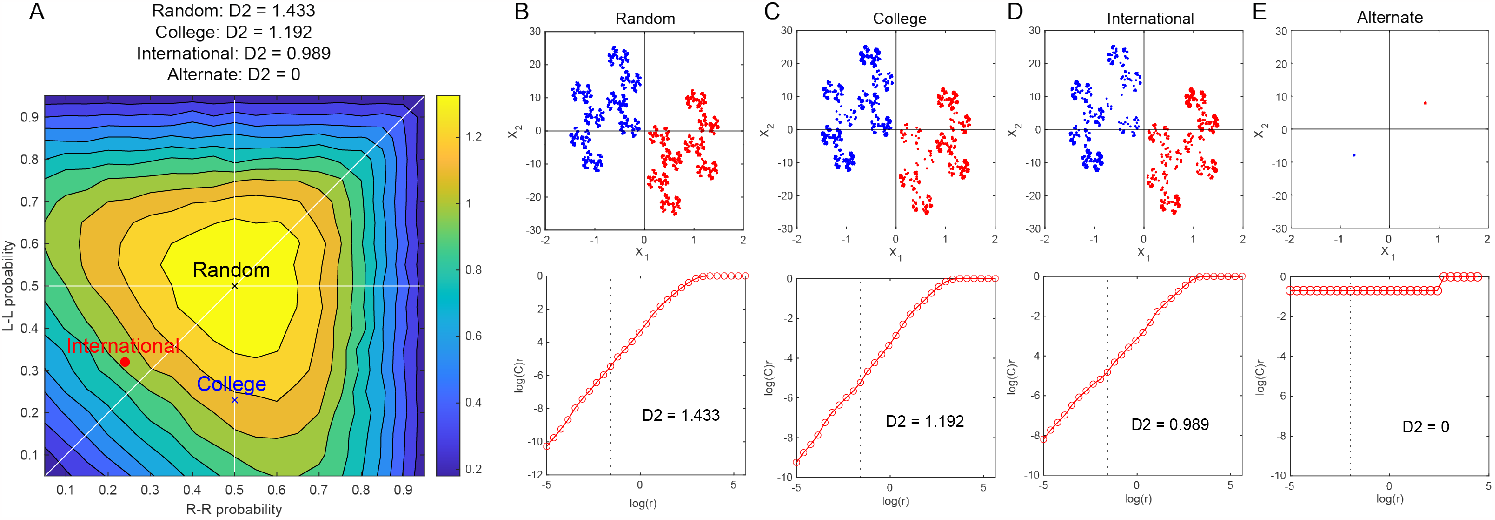
Correlation dimensions on Poincaré maps were obtained through simulation experiments, corresponding to various second-order Additionally, Poincaré maps and correlation dimensions were obtained for four representative state transition probabilities (B-E).**Figure 5—figure supplement 1**. State transition probabilities for the simulation experiment and examples of results. A to D show the second-order state transition probabilities for the pseudo-international, alternate, and random series, respectively. E to H show the examples of the results of the simulation experiments corresponding to A to D, respectively. The black step waves show the external input pattern; red and blue lines show the output pattern for *x*_1_; and red and blue dotted lines show the output pattern for *x*_2_. **Figure 5—figure supplement 2**. Correlation dimensions in real matches compared to pseudo-international and pseudo-collegiate sequences.

The pseudo-international and pseudo-collegiate sequences, which were generated based on the second-order state transition probabilities derived from the sequences of both international and collegiate matches, were located between complete regularity and complete randomness, as shown in ***Figure 5*** A. The Poincaré maps for the college and international levels corresponded to ***Figure 5*** C and D, respectively, with correlation dimensions of 1.192 and 0.989, respectively. This indicates that the pseudo-collegiate sequence is closer to a completely random sequence than to the pseudo-international sequence. Compared with real matches, the correlation dimensions for international and collegiate matches were similar to the results obtained from the pseudo-sequences. The correlation dimension for international matches was smaller than that for collegiate matches, which is consistent with the simulation results (***Figure 5—figure Supplement 2***).

## Discussion

Interpersonal competition in court-net sports requires successive responses to an opponent’s shot, with quick decision-making and appropriate execution within short-time intervals. Because one cannot completely predict others’ intentions, court-net sports serve as an ideal platform for observing human characteristics as in a circular argument. Specifically, court-net sports helps us observe how individuals who attempt to control unpredictable others in sequential and reciprocal situations strive to minimize uncertainties in an environment. To this end, we applied the SHDS model (***Nishikawa and Gohara, 2008a***,b; ***Yamamoto et al., 2018***) to analyze shot angles and on-court movements in international and collegiate soft tennis matches. Furthermore, by incorporating the mass-spring-damper model into the SHDS, we conducted simulation experiments to investigate the fractal nature of on-court movements resulting from the switching of input sequences. This analysis helped elucidate the characteristics of regularity in the sequence of shot angles observed in actual matches.

One of the main findings of this study is that using regularity to reduce the uncertainty of unpredictable intentions of others is an essential interpersonal skill in controlling others in sequential and symmetric reciprocal competitive situations. This strategy is believed to involve skilled players capitalizing on their opponent’s intention to predict or reduce uncertainty (***Friston, 2010; Friston et al., 2017***). They create their regular patterns to make it easier for the opponent to predict and then disrupt that regularity to confound the opponent’s predictions. This is evident from the observed skilled players’ regularity in the shot angle, which falls between a completely random series and a consistently alternating left-right series. This is also apparent in ***Figure 2—figure Supplement 1*** and ***Figure 2—figure Supplement 2***, where regularity is not used in the final phase of the rally but predominantly before it. Conversely, collegiate players exhibit less regularity and tend to approach randomness in their actions, which may be attributed to their efforts to avoid their opponents’ predictability (***Glover and Dixon, 2017***). These results suggest that skilled players possess interpersonal motor skills that involve initially behaving consistently to reduce others’ uncertainty, and then prompting others to predict the regularity of their actions as a means of deception. To achieve this, the motor skill is used to create regular patterns in the opponents’ predictions. In other words, the players’ strategy is not that of preventing the opponents from predicting them from the beginning but that of deceiving the opponents by initially prompting them to predict the players’ behaviors (***Vesper et al., 2010, 2011; Sabu et al., 2020***). This action decreases the unpredictability of the opponent and increases the predictability of the opponent’s behavior. Both individuals want to avoid surprising themselves in their actions but face a dilemma as they want to surprise others. It argues that sequential, symmetric, reciprocal, and competitive interactions form a circular argument.

We also found that in actual matches, skilled players exhibited more regularity in their shot angle series than collegiate players. We also observed a third-order sequence effect, namely hysteresis, in the on-court movements of opposing players, which correlated with the regularity of the series. Previous well-controlled experiments demonstrated a third-order sequence effect, known as hysteresis, in tennis hitting behavior in response to random input switching (***Yamamoto and Gohara, 2000***). This implies that even when hitting the same forehand, the current hitting pattern will differ depending on whether the preceding hit is the forehand or backhand. Hysteresis corresponds to the temporal development of the Cantor set with rotation, which has been posited as a fractal transition in the SHDS (***Gohara and Okuyama, 1999a; Nishikawa and Gohara, 2008a***,b). This hysteresis is believed to stem from the inertia in the rotational motion of hitting, causing a time lag in the rotation of the trunk. In the current experiment, we observed the same hysteresis as in the controlled experiment for the movements involved in hitting the ball in response to the regularity of the opponent’s shots during rallies in real matches. This observation may also be attributed to a delay in returning to the home position after taking a shot in court. This is commonly referred to as a disruption of the opponent’s system.

Models of coupled oscillators, such as the HKB models (***Haken et al., 1985***), have explained the synchronization between the two systems (***Schmidt et al., 1990; Kijima et al., 2012; Okumura et al., 2012; Nalepka et al., 2019***). However, because these models represent autonomous systems, they do not consider the individual behaviors of the two systems. In contrast, SHDS is a model of a non-autonomous system that explicitly incorporates temporal external inputs, allowing us to analyze individual players’ behaviors. Although previous applications of SHDS to human movements have experimentally controlled external temporal inputs (***Yamamoto and Gohara, 2000; Suzuki and Yamamoto, 2015; Hirakawa et al., 2016, 2017***), the present results were obtained during an actual match between two players without controlling for external inputs. Thus, incorporating this external temporal input into SHDS may enable us to elucidate the underlying principles of seemingly complex behaviors in human interactions.

This study showed that two players’ behavior in court net sports can be described as two-coupled SHDS and that expert players use interpersonal strategies that decrease the unpredictability of the opponent through regularity in their own actions. This finding regarding interpersonal strategy could help policymakers in the field of business and organizational behavior formulate policies that facilitate their competitiveness.

## Methods and Materials

### Data acquisition in real matches

We recorded nine male singles in the 8th Asian Soft Tennis Championship in Chiba, Japan, in November 2016, and nine male singles in the second division of the Tokai collegiate league in Tsu, Japan, in September 2017, using a video camera (SONY, HDR-CX630, 30 fps). The international matches and collegiate matches scored 353 and 300 points, respectively, in all nine matches. However, 39 and 20 points of international and collegiate matches, respectively, were excluded from the analysis because of recording problems.

The players in international matches represented the top three countries in the world, Chinese Taipei, Republic of Korea and Japan. The players in collegiate matches played in the second division of the local (tokai) regional collegiate league.

The ground level of the center of the feet of both players was digitized at a sampling rate of 30 Hz from service to the end of the rally, and two-dimensional Cartesian coordinates were reconstructed using the direct linear transformation method(***Abdel-Aziz and Karara, 1971***). The origin was set at the center of the court; the X-axis was parallel to the baseline; and the Y-axis was parallel to the side line. All positional data were low-pass filtered using a dual-pass zero-lag fourth-order Butterworth filter with a cut-off frequency of 1 Hz. Further, the moments at which the ball hit were identified through a frame-by-frame observation.

The rallies over nine shots, including the service, were extracted from all points because we analyzed more than four successive shots of each player to examine the regularity of the return map and third-order sequence effect of hysteresis in the continuous movements. As a result, 101 and 31 points were analyzed for the international and collegiate matches, respectively, and the ratio of the analyzed data was 28.6 % and 10.3 % of all points, respectively.

All participants provided written informed consent prior to recording. The procedures were approved by the Internal Review Board of the Research Center of Health, Fitness, and Sports at Nagoya University and conformed to the principles of the Declaration of Helsinki. Approval from the Japan Soft Tennis Association was obtained for the use of videos in this study.

### Analysis of the shot sequences in a rally

#### State variables

To depict the successive decision making using the return map as a discrete dynamical system, we defined the state variables which were the shot course as state variables in decision making. For the continuous dynamical system, continuous movements as trajectories in the cylindrical phase space were defined as two state variables—the position and velocity of the movements.

The shot course consisted of the shot angle Θ and shot length ΔL as candidate state variables. The shot angle Θ is calculated as the inner angle between each vector of the opponent’s shot trajectory and the subsequent shot trajectory. This variable can be calculated for shots, including the service return to the last shot in each rally. We defined the angular deviation of the shot trajectory to the right of the vector inverse to the preceding shot trajectory as positive and that to the left as negative (***Figure 6*** A). Shot length ΔL is defined as the depth of a player’s shot relative to the position at which the opponent released the preceding shot. If the opponent is forced to move backward by the player’s shot, ΔL is calculated as positive (***Figure 6*** B).

**Figure 6.**
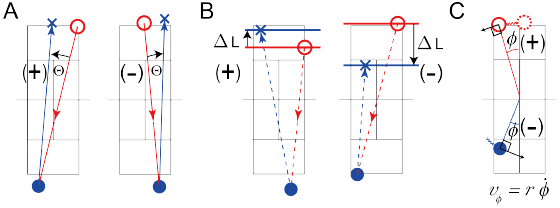
Definition of state variables. A and B show the definition of shot angle (Θ) and shot length (ΔL) for the discrete dynamical system, respectively. In A and B, red circles show the opponent’s hitting positions; red lines show the preceding shot; blue circles show the current hitting position; blue lines show the current shot; and blue crosses show the opponent’s next hitting position. C shows position (*ϕ*) and velocity (*v*_*ϕ*_) defined on polar coordinates with the origin at the center of the court.

Players’ positions and velocities were defined in polar coordinates by the origin at the center of the court. The position is defined as the polar angle *ϕ*, and the velocity is defined as the tangential velocity *v*_*ϕ*_ = *rϕ* (***Figure 6*** C). The deuce court side is defined as positive and the advantage side as negative.

These four variables, Θ, ΔL, *ϕ* and *v*_*ϕ*_ were calculated for sequences that included more than nine shots (the reason for this limitation on the length of the rally is explained in the following paragraph). Each of the four variables was calculated for 202 sequences (101 rallies × two players) and 62 sequences (31 rallies × two players) for international and collegiate matches, respectively.

#### Return map analysis

The discrete dynamics of the two variables, Θ and ΔL, can be analyzed on a return map, similar to a Lorenz map (***Lorenz, 1963***). The state of each data point for a given player Θ_*n*+1_ can be visualized by plotting it as a vector relative to that of *n*^*th*^ previous shot Θ_*n*_. The transition pattern of the points represents the trajectory of the state in a discrete dynamical system.

The return map analysis can be applied to the sequential data {Θ} = Θ_1_, Θ_2_…Θ_*n*_ as shown in ***Figure 2*** A. Data set of the first two points are represented by the point (Θ_1_, Θ_2_), indicated by the filled marker, and move to the next point (Θ_2_, Θ_3_) through the point (Θ_2_, Θ_2_) as shown in ***Figure 2*** E. Therefore, the data set of the last two points can be represented by the point (Θ_*n*−1_, Θ_*n*_) following the point (Θ_*n*−1_,Θ_*n*−1_). As can be seen in ***Figure 2*** A and B, if the sequence fluctuates periodically, the points on the map move along the rotated trajectory in ***Figure 2*** E and F, whereas the sequence fluctuates and moves to the fixed point in ***Figure 2*** C and D, and the points move from the point asymptotically, as shown in ***Figure 2*** G and H. Thus, we can determine the type of fluctuation in a given sequence from the pattern of the moving trajectory in the return map space.

We postulated that each of the two variables, Θ and ΔL, fluctuates in either of these two patterns periodically or asymptotically and that the pattern of the given variable can be mathematically determined by analyzing the relationship between the geometrical distribution of Θ and the linear function *X*_*n*+1_ = *a X*_*n*_ + *b*. For example, if the distribution pattern of Θ on the map fits the linear regression function *X*_*n*+1_ = *a X*_*n*_ + *b*, this relationship can be easily determined by confirming the parameter *a* (***Garfinkel et al., 1992; Schiff et al., 1994***). When *a* < 0, the point on the map moves along the rotational trajectory in ***Figure 2*** E and F, whereas Θ_*n*_ alternately decreases or increases, as shown in ***Figure 2*** A and B. By contrast, in the case of 0 < *a*, the trajectories are asymptotically close to, or away from, a certain point in ***Figure 2*** G and H, whereas Θ decreases or increases gradually to that point, as shown in ***Figure 2*** C and D, respectively. Additionally, if |*a*| < 1 the points are attracted to the fixed point (i.e., an “attractor”) as shown in ***Figure 2*** F and H, whereas in the case of |*a*| > 1, the points move away from a given point (i.e., a “repeller”) as shown in ***Figure 2*** E and G.

If the rally includes ten shots and starts with a service shot by player X and ends with a shot by player Y, player X and Y has four and five successive data points of Θ and ΔL, respectively, which were calculated from the shot angles excluding that of the service (his/her 1st shot) by player X. These data points calculated from the shot angles of players X and Y can be projected onto the corresponding three and four points on the return map, respectively. Then, for the data shot by player X, the linear fitting analysis can be applied to only one sequence of three successive points on the return map calculated from second to fifth shots of player X. On the other hand, for player Y, four successive plotted point fitting can be applied to two sequences of three successive points on the map representing the first to forth shots and that representing the second to fifth shots, and the four-point fitting can be applied to successive data points from the first to fifth shots.

Thus, *N* + 1 successive shots are required to calculate *N* points on the return map. Therefore, a rally must include at least eight shots in total to order to project at least three points on the return map. In this case, the receiver returns at least four shots, which are sufficient to calculate the three points on the map. However, for the shot sequence of the server, only two points can be projected on the map calculated from three shots, excluding the service shot. Thus, to determine the linear function on the return map, the length of the shot sequence must be greater than four, that is, nine shots in total. Therefore, the range of lengths of the shot sequences was set to four, five, six, seven, and eight to calculate three, four, five, six, and seven points on the return map, respectively. Further, as explained above, we can extract *i* + 1 sequences of *N* successive shots from the rally, including *N* + *i* return shots, for each player. We independently analyzed each *i* + 1-sequence on the return map extracted from the same rally.

#### Statistical procedure of linear fit analysis

To determine the function that significantly fits the given data, a linear fitting analysis was applied from three to seven plotted points in each rally. The point on the return map was calculated for each data sequence with either of the five lengths (three, four, five, six, and seven points on the return map), and a linear fit analysis was applied to all these sequences of points on the return map. The significance of fitting to the linear function *X*_*n*+1_ = *a X*_*n*_ + *b* was estimated using the coefficient of determination *R*^2^ and the significance of the regression model, and the significance level was set to *R*^2^ > 0.8 or *p* < .05. This is because the coefficient of determination, *R*^2^ decreases for longer fitting points, while the probability of significance of the regression model increases for longer fitting points. The ratio of the number of significantly fitted sequences of a given length to the total number of sequences of the same length observed in the same match was calculated. This ratio was calculated independently for each sequence with five lengths for each of the nine matches performed by international and collegiate players.

The significance of these ratios was tested through a comparison of the ratios calculated from the 200 surrogate data points obtained from random shuffling data with a given length of the shot sequence(***Theiler et al., 1992***) (***Figure 1—figure Supplement 1***). These surrogate data were also fitted to a linear function using the same procedure and criteria that were applied to the points calculated from the actual data. The ratio of the number of significantly fitted data points to all 200 surrogate data points was calculated independently for each sequence, with five lengths for each of the nine matches performed by international and collegiate players.

The significance of the difference in the ratio between the actual and surrogate data was then estimated by testing the equality of the given proportion hypothesis independently for each skill level and length of successive shots. The level of significance was set at p < 0.05. This significant difference indicates that such regularity is hidden in the sequence as a rotationally or asymptotically moving attractor or repeller.

#### Analysis of trajectories in cylindrical phase space

To examine a continuous dynamical system, the position (*ϕ*) and velocity (*v*_*ϕ*_) are designated *x*_1_ and *x*_2_, respectively. The output pattern— the movements of players—was depicted as trajectories in the hyper-cylindrical phase space ℳ, which started at the moment of the opponent’s preceding shot (i.e., the Poincaré section Σ(*θ* = 0)), until the next opponent shot (Σ(*θ* = 2*π*)). The trajectory around the cylinder represents a single cycle.

The analyzed data were the output patterns corresponding to a well-fitted input pattern on the return map analysis, i.e., the sequence of the opponent’s shot angles. Further, to examine the hysteresis of the output pattern corresponding to the input sequence, we considered the third-order sequence effect of the input. In this study, two input patterns were used: right (R) and left (L). Thus, the third-order sequence effect was 2^2^ = 8 pattern, that is, RRR, LRR, LLR, RLR, LLL, RLL, RRL, and LRL. The well-fitted input pattern consisted of four-to-eight sequences of shots; for the four-shot sequence, we could only observe two series of third-order sequence effects–from the first to the third shot and from the second to the forth shot.

We also calculated the second-order transition probability for the well-fitted input sequences— RR, LR, LL, and RL, because if we observed the second-order sequence effect on the Poincaré section Σ, it would indicate eight types of trajectories in the cylindrical phase space(***Yamamoto and Gohara, 2000***). To examine the second-order sequence effect on the Poincaré section Σ, a MANOVA using Wilks’ Λ of position *ϕ* and velocity *v*_*ϕ*_ as multivariate variables on five Poincaré sections Σ(*θ* = 0, *π/*2, *π*, 3*π/*2, 2*π*) and the effect size 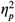 were calculated.

### Simulation experiment

Model for continuous movement on the court corresponding to external input First, the damped mass-spring model, which was applied to discrete movements in the shepherding task (***Nalepka et al., 2019; Patil et al., 2020***), was applied to the movements of players on the court.

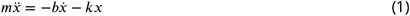

where *x* represents the position of an object or body, 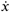 the velocity of *x* over time, and 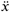 the acceleration of *x* over time. Further, *m* is the object mass, *b* is a damping parameter that resists motion (for *b* > 0), and *k* is a spring force or stiffness parameter that induces motion (for *k* > 0). When *b* = 0, this mass-spring model is termed a simple harmonic oscillator. If we set *m* = 1 for normalization, ***Equation 1*** can be transformed into

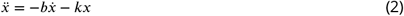

To apply this damped mass-spring model to the switching dynamics, this equation can be expressed as its modification into the first-order differential equation for a dynamical system. We substitute 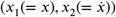 with *x*, which is generally expressed by the following equation (***Gohara and Okuyama, 1999a; Gohara et al., 2000***):

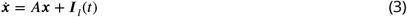

***Equation 2*** is expressed as follows:

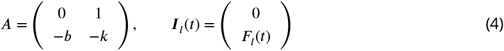

where *F*_*l*_(*t*) is a series of two types of external inputs, [1, −1, −1, −1, 1, …], with constant values. *I*_*l*_(*t*) switches with time interval *T*.

#### Series of external temporal inputs

We defined the four types of 2,000 series of external inputs to examine the behavior of the system. First, a pseudo-international and a pseudo-collegiate series are generated based on the second-order state transition probabilities observed in international and collegiate matches (***Figure 3*** A, ***Figure 5—figure Supplement 1*** A and B). Second, a periodic series is alternately generated on the right and left sides (***Figure 5—figure Supplement 1*** C). Finally, a random series is generated based on a uniform distribution from 0 to 1 (***Figure 5—figure Supplement 1*** D).

#### Parameter tuning

We chose *b* and *k* in ***Equation 2*** and the time interval *T* to switch to the external input so that the switching movement pattern could be observed (***Figure 5—figure Supplement 1*** E-H). Thus, we set *b* = 10, *k* = 500 in ***Equation 2***, and the time interval was set to 100 ms. However, this time interval did not correspond to the actual time, but only for the calculation.

## Supporting information

Readme.txt

International match data

Collegiate match data

## Acknowledgments

We thank Riko Kudo for her preliminary data analysis and Koji Kadota for valuable comments on the manuscript.

## Appendix 1 Switching hybrid dynamical system

First, to consider the switching movement pattern corresponding to the external temporal input *I*(*t*), a nonautonomous system (a system that depends on the external input) is defined using ordinary differential equations of the type:

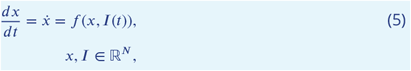

where *x* and *f* represent the state and vector fields, respectively(***Gohara and Okuyama, 1999a***). When the input *I*(*t*) varies slowly compared to state *x*, the input can be regarded as a constant, resulting in a bifurcation parameter. In human movements, finger or wrist movements correspond to state *x* and regular changes in rhythm are equivalent to the input *I*(*t*)(***Kelso, 1984; Haken et al., 1985***). Conversely, when a change in input *I*(*t*) cannot be neglected compared to the change in state *x*, the way the vector fields change with time should be considered. In this case, the input is introduced as a set 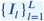 of temporal input *I*_*l*_ with finite duration *T*_*l*_. For example, for hitting movements in tennis, a temporal input with a finite duration corresponds to the forehand or backhand of the opponent’s shot. The hitting movement pattern, state *x*, switches between the forehand and backhand strokes corresponding to the opponent’s shot course. Each input element is defined as the period of a periodic time function. The input corresponds to a set of sampling points in the parameterized space of the function. For each sampling point, a continuous dynamical system is defined by the vector field *f*_*l*_ in hypercylindrical phase space ℳ. A discrete dynamical system is defined by the iterative function *g*_*l*_ in the global Poincaré section Σ, which is a subspace of ℳ.

When the same input *I*_*l*_ is repeatedly fed into the system, the trajectory, *γ*_*l*_ converges onto an attractor in space ℳ(***Gohara and Okuyama, 1999a***). This attractor was designated an excited attractor to emphasize that it was excited by an external input. Excited attractors represent movement patterns such as forehand or backhand strokes, corresponding to a ball course in tennis. Generally, when external inputs are switched stochastically, the trajectory cannot converge to either excited attractor. However, the following results were demonstrated both analytically and numerically(***Gohara and Okuyama, 1999b; Yamamoto and Gohara, 2000***). Trajectory *γ*_*l*_(*C*) in the hypercylindrical space ℳ converges to set Γ(*C*) starting from the initial set *C* in the Poincaré section Σ. Two sets, Γ(*C*) and *C*, are attractive and invariant sets with fractal-like structures that satisfy the following equations:

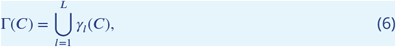

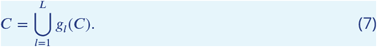

The trajectory set Γ(*C*) corresponding to the input set 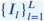 is obtained from the union of the trajectory set *γ*_*l*_(*C*) with each input *I*_*l*_ and is composed of transient trajectories between the excited attractors. The initial set *C* in the Poincaré section Σ is obtained from the union of sets 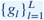 of the iterated functions *g*_*l*_ in the Poincaré section (***Hutchinson, 1981; Barnsley, 1993***). Because the boundaries between these transient trajectories are fractal, we can refer to the fractal transitions between the two attractors(***Gohara and Okuyama, 1999b***).

The proposed integrated model was a switching hybrid dynamical system(***Nishikawa and Gohara, 2008a***,b). Here, we assume a system with higher and lower modules that interact with each other by a switching input *I*_*l*_(*t*) from the higher to the lower module and by a feedback signal *x*(*t*) from the lower to the higher module. Further, an external input *I*_*ext*_(*t*) is fed into the higher module as shown in ***Appendix 1–Figure 1***. Here, the higher module corresponds to the brain or prefrontal cortex as a discrete dynamical system(***Miller, 2000; Sakagami and Tsutsui, 1999***), where the higher module transforms both the external input pattern *I*_*ext*_(*t*) and feedback signal *x*(*t*) into the input pattern *I*_*l*_(*t*). The lower module corresponds to the human motor system as a continuous dynamical system, described by Eq. 5.

**Appendix 1—figure 1.**
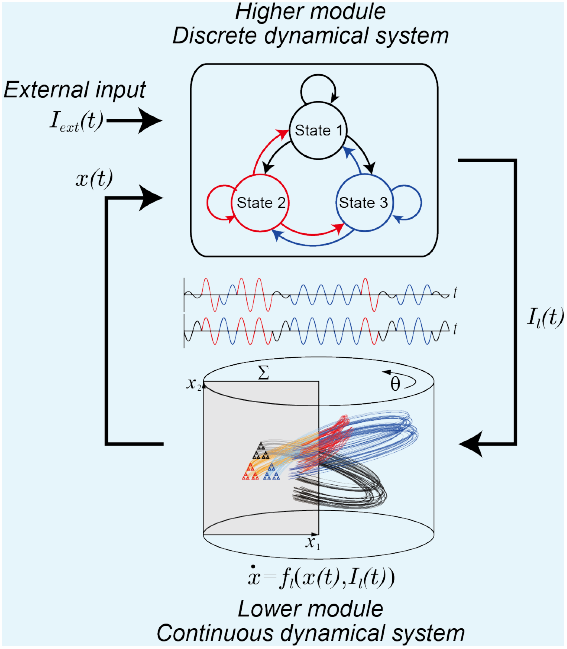
Schematic representation of switching hybrid dynamics composed of a discrete dynamical system as the higher module and a continuous dynamical system as the lower module with a feedback loop. This system is non-autonomous.

## Appendix 2 Skill difference in the rally length

A histogram indicating the distribution of the number of shots per rally was calculated for each of the nine matches for each skill level ***Appendix 2–Figure 1***. To test the significance of the group difference in the number of shots included in each rally, a repeated-measures ANOVA with Greenhouse-Geisser correction for level (2) by the number of shots (20) was applied to the histogram. Because rallies with more than 20 shots were extremely rare, those frequencies were included in the frequency of 20 shots. The Holm method (***Holm, 1979***) was used for multiple comparisons. In addition, the chi-square test and residual analysis were applied to the frequencies of winners and errors for fewer than nine successive shots and over nine shots for both levels. These statistics were performed using programs written using R software.

The means and standard deviations of the rally lengths were 8.1±6.53 and 4.7±3.19 in international and collegiate matches. The two-way repeated-measures ANOVA of levels × the rally length revealed that each main effect of skill level and rally length was significant (skill levels: *F* (1, 16) = 13.67, *p* = .002, 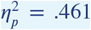; rally length: *F* (7.47, 119.48) = 93.38, *p* = .000, 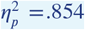), and the effect of the interaction of the factors was also significant (*F* (7.47, 119.48) = 2.45, *p* = .020, 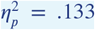). The simple main effect of skill level was significant for each frequency of rallies with three, nine, ten, and over 20 shots. Relatively short rallies, such as those that include only three successive shots(***Fitzpatrick et al., 2019***), occur more frequently in collegiate matches than international ones. However, longer rallies including nine, ten, or over 20 successive shots, occurred more frequently in international matches than in collegiate matches.

The cumulative frequencies of shots were 2,529 and 1,325 in international and collegiate matches, respectively. More than nine shots occurred in 63 % of international matches and 27 % of collegiate matches.

**Appendix 2—figure 1.**
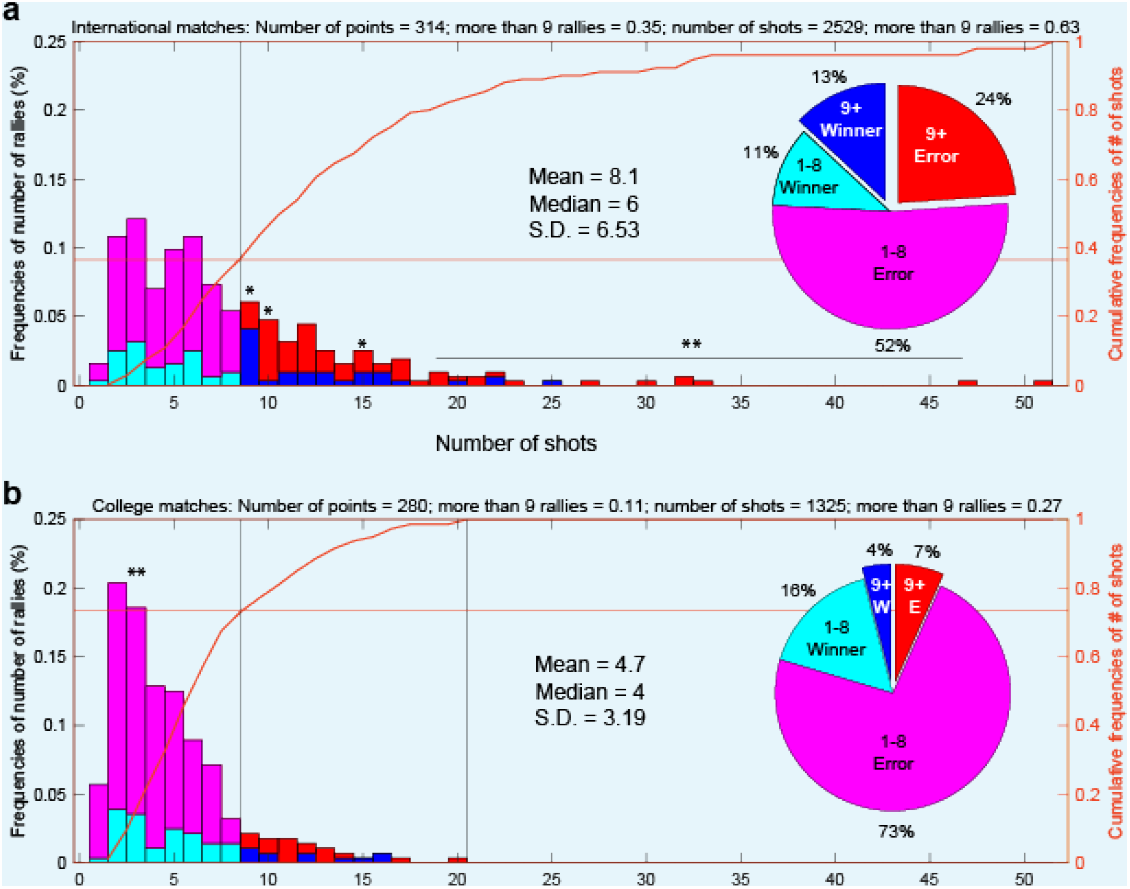
Frequencies of number of rallies and ratios of winners and errors. (**a**) and (**b**) show international and college matches, respectively. The blue bars show the frequency of winners in rallies with over nine shots, the cyan bars show the frequency of winners in rallies with fewer than nine shots, the red bars show the frequency of errors in rallies with over nine shots, and the magenta bars show the frequency of errors in rallies with fewer than nine shots. Orange curves show the cumulative frequencies of the number of shots (right axis). Pie charts showing the proportions of winners and errors for all points.

---

**Figure 1—figure supplement 1.**
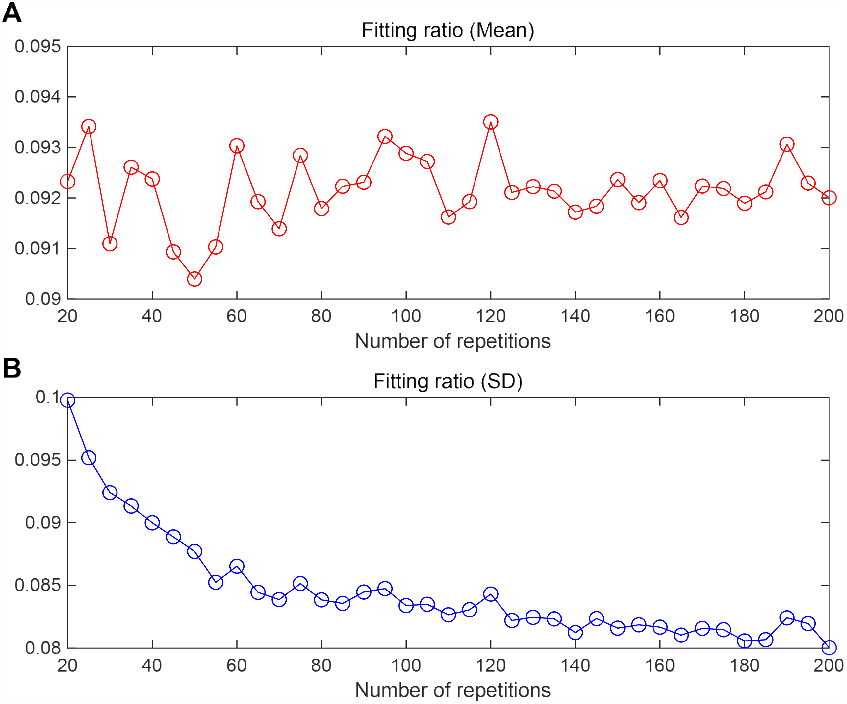
Means and standard deviations (SDs) of the fitting ratio by the number of random shuffling repetitions for surrogate data. A shows the change in the mean of the fitting ratio for 20—200 repetitions. B shows the change in SD of the fitting ratio for 20—200 repetitions.

**Figure 2—figure supplement 1.**
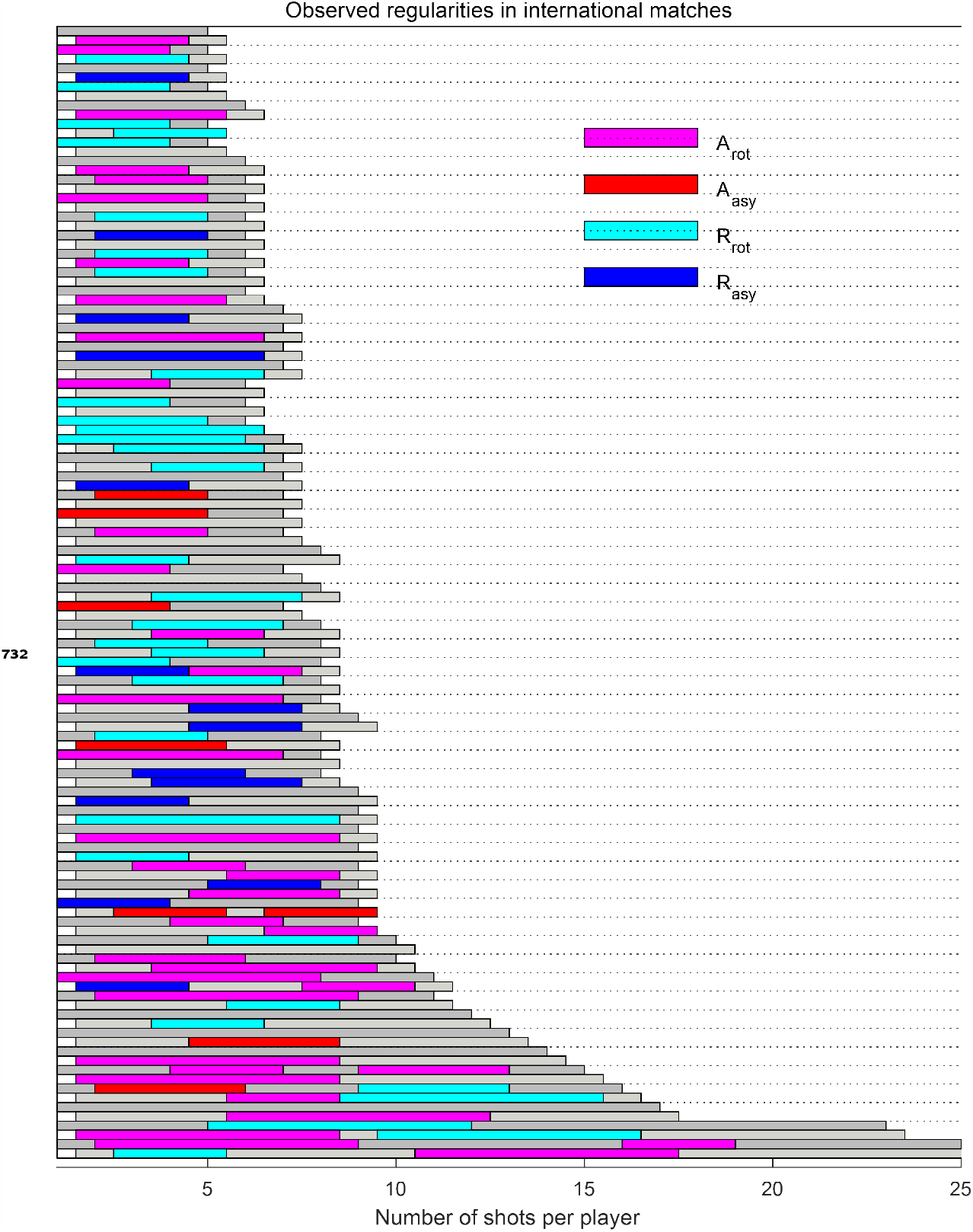
Observed regularities in each rally per player in international matches

**Figure 2—figure supplement 2.**
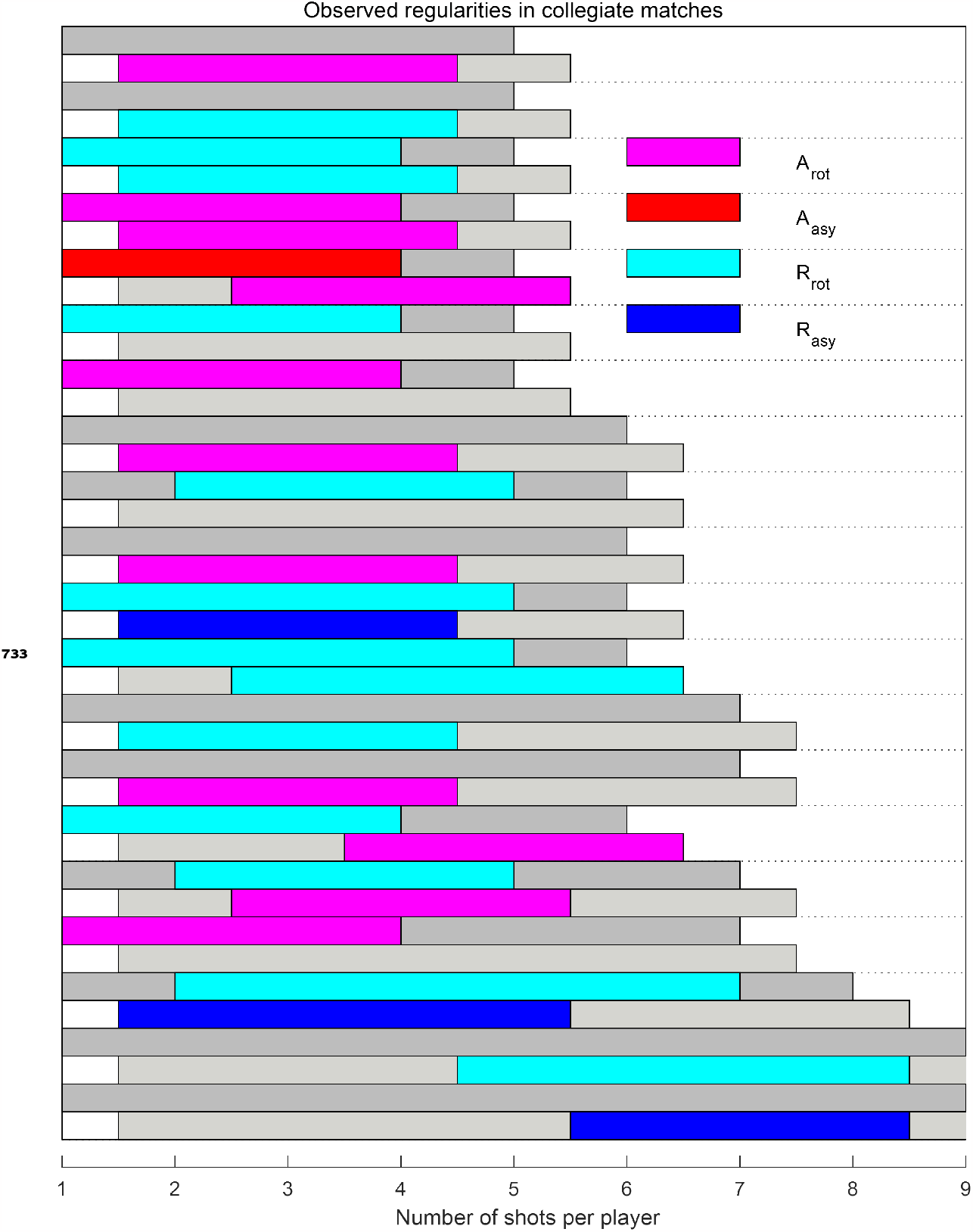
Observed regularities in each rally per player in collegiate matches

**Figure 3—figure supplement 1.**
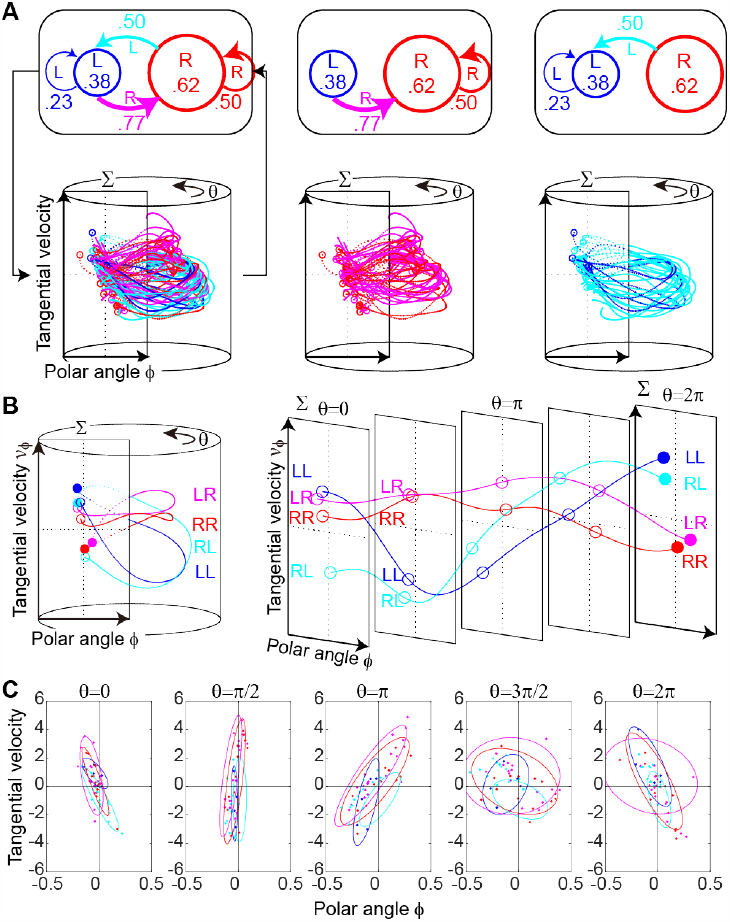
Switching hybrid dynamics in the collegiate match. A-C shows the same in ***Figure 3***.

**Figure 3—figure supplement 2.**
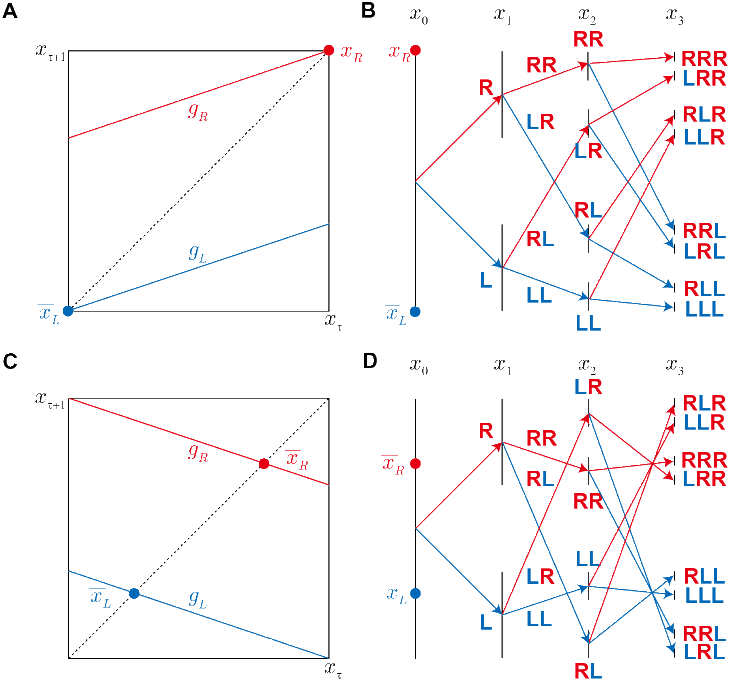
Hierarchical fractal structure. This demonstrates how the Cantor set can be constructed using two iterative functions. A and C show the iterative function *g*_*R*_ and *g*_*L*_ transform state *x*_*τ*_ to the next state *x*_*τ*+1_. In A, the lower-left and upper-right corners are the attractive fixed points 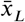 for *g*_*L*_ and 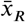 for *g*_*R*_, respectively. B and D show the hierarchical structure of the fractal corresponding to the sequence of right (R) and left (L) side inputs from *x*_0_ to *x*_3_. C and D are the same as A and B, except that the transformations of iterative function *g*_*R*_ and *g*_*L*_ are rotated around the fixed points 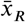 and 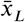, respectively. We named this the Cantor set with rotation.

**Figure 3—figure supplement 3.**
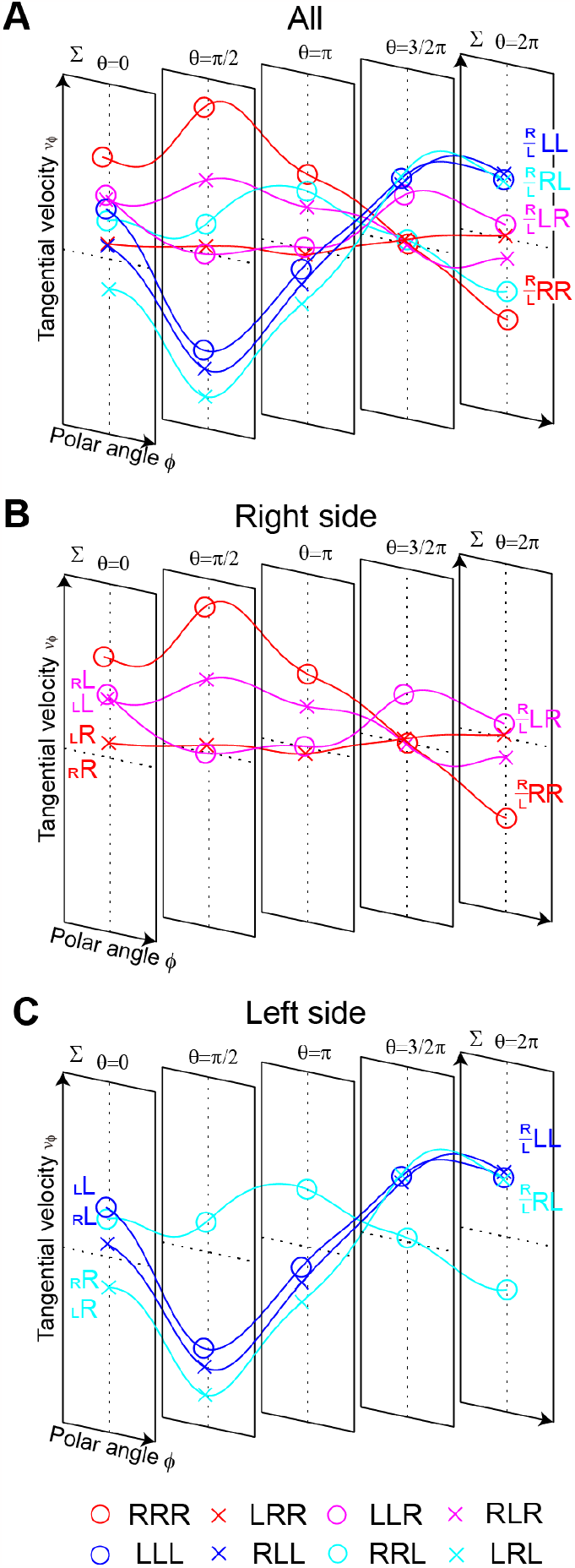
Eight trajectories of transitions between two excited attractors. The mean trajectories are expanded into the two-dimensional plane, using *θ* −(*x*_1_, *x*_2_) as the switching input. The trajectories are drawn with the third-order sequence effect. A shows eight mean trajectories. B and C show four mean trajectories corresponding to the right and left side inputs, respectively.

**Figure 4—figure supplement 1.**
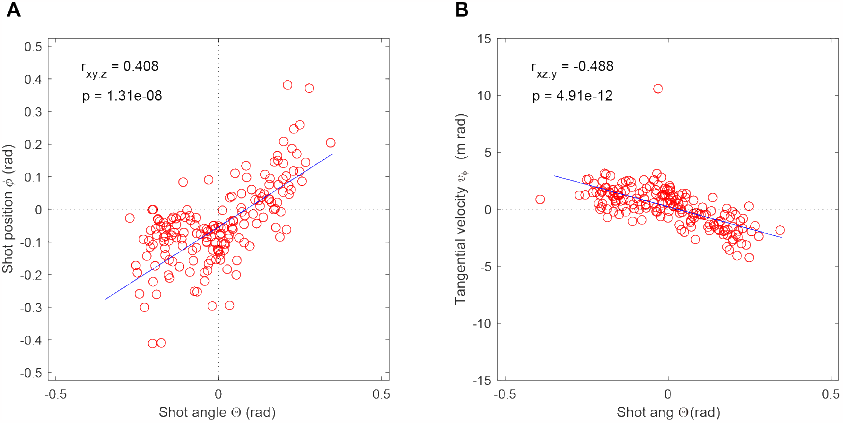
Partial correlation. A shows the partial correlation between the shot angle Θ and polar angle *ϕ*. B shows the partial correlation between the shot angle Θ and tangential velocity *v*_*ϕ*_.

**Figure 5—figure supplement 1.**
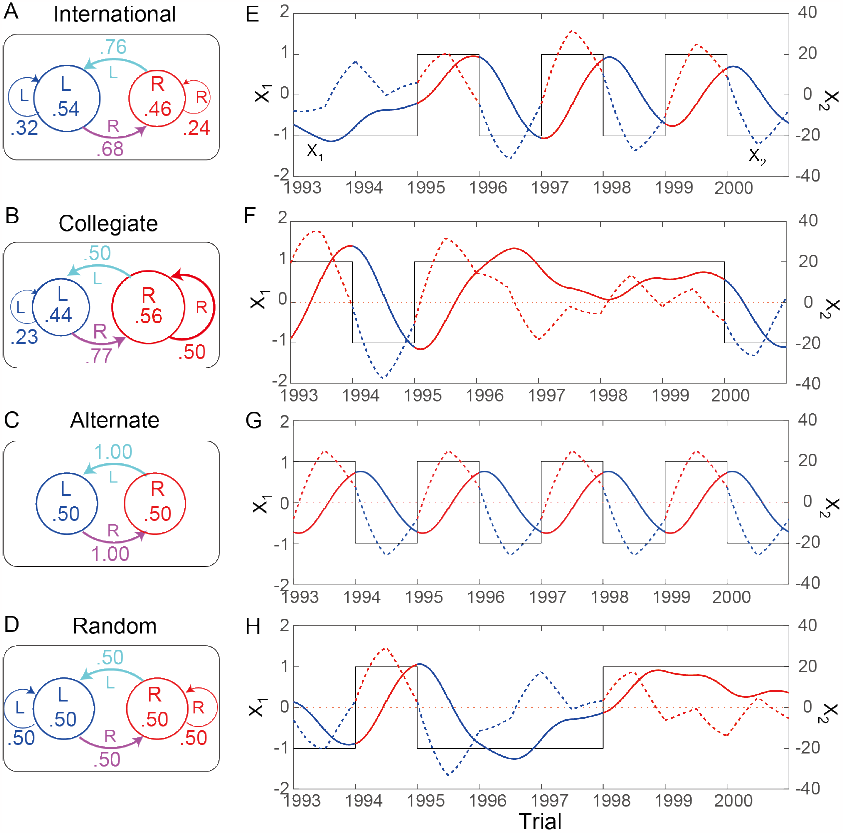
State transition probabilities for the simulation experiment and examples of results. A to D show the second-order state transition probabilities for the pseudo-international, alternate, and random series, respectively. E to H show the examples of the results of the simulation experiments corresponding to A to D, respectively. The black step waves show the external input pattern; red and blue lines show the output pattern for *x*_1_; and red and blue dotted lines show the output pattern for *x*_2_.

**Figure 5—figure supplement 2.**
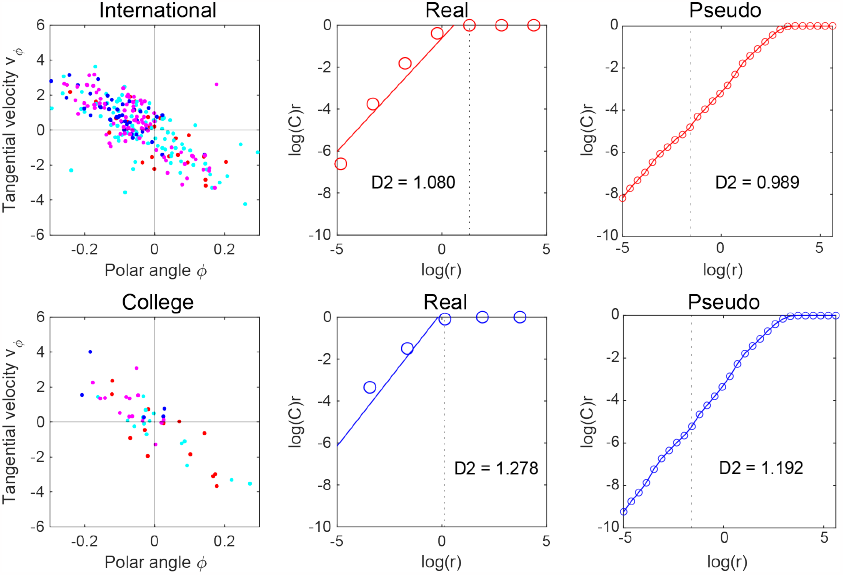
Correlation dimensions in real matches compared to pseudo-international and pseudo-collegiate sequences.

