## Supplementary material for "How to control unpredictable others in court-net sports": Readme.txt

International.zip and Collegiate.zip includes raw positional data.
Each file name shows categories (n/u), match ID, game count, point.
For example, n1_2_1.txt means the rally of international match, match ID=1, second game, and 1st point.

Data columns show following Cartesian coordinates.
X_a, Y_a, X_b, Y_b, Impact_X_a, Impact_Y_a, Impact_X_b, Impact_Y_b
Data sampling rate was 30 Hz.
